# A cytochrome *bd* repressed by a MarR family regulator confers resistance to metals, nitric oxide, sulfide, and cyanide in *Chromobacterium violaceum*

**DOI:** 10.1101/2024.08.06.606881

**Authors:** Bianca B. Batista, W. Ryan Will, Vinicius M. de Lima, Ferric C. Fang, José F. da Silva Neto

## Abstract

*Chromobacterium violaceum* is a ubiquitous environmental pathogen. Despite its remarkable adaptability, little is known about the mechanisms of stress resistance in this bacterium. Here, in a screen for iron-susceptible transposon mutants, we identified a cytochrome *bd* that protects *C. violaceum* against multiple stresses. The two subunits of this cytochrome *bd* (CioAB) are encoded by the *cioRAB* operon, which also encodes a GbsR-type MarR family transcription factor (CioR). A Δ*cioAB* mutant strain was sensitive to iron and the iron-requiring antibiotic streptonigrin and showed a decrease in siderophore production. Growth curves and survival assays revealed that the Δ*cioAB* strain was also sensitive to zinc, hydrogen peroxide, nitric oxide, sulfide, and cyanide. Expression analysis showed that the promoter activity of the *cioRAB* operon and the transcript levels of the *cioAB* genes were increased in a Δ*cioR* mutant. CioR bound the promoter region of the *cio* operon *in vitro*, indicating that CioR is a direct repressor of its own operon. Expression of the *cio* operon increased at high cell density and was dependent on the quorum-sensing regulator CviR. As cyanide is also a signal for *cio* expression, and production of endogenous cyanide is known to be a quorum sensing-regulated trait in *C. violaceum*, we suggest that CioAB is a cyanide-insensitive terminal oxidase that allow respiration under cyanogenic growth conditions. Our findings indicate that the cytochrome *bd* CioAB protects *C. violaceum* against multiple stress agents that are potentially produced endogenously or during interactions with a host.

**IMPORTANCE:** The terminal oxidases of bacterial respiratory chains rely on heme-copper (heme-copper oxidases) or heme (cytochrome *bd*) to catalyze reduction of molecular oxygen to water. *Chromobacterium violaceum* is a facultative anaerobic bacterium that uses oxygen and other electron acceptors for respiration under conditions of varying oxygen availability. The *C. violaceum* genome encodes multiple respiratory terminal oxidases, but their role and regulation remain unexplored. Here, we demonstrate that CioAB, the single cytochrome *bd* from *C. violaceum*, protects this bacterium against multiple stressors that are inhibitors of heme-copper oxidases, including nitric oxide, sulfide, and cyanide. CioAB also confers *C. violaceum* resistance to iron, zinc, and hydrogen peroxide. This cytochrome *bd* is encoded by the *cioRAB* operon, which is under direct repression by the MarR-type regulator CioR. In addition, the *cioRAB* operon responds to quorum sensing and to cyanide, suggesting a protective mechanism of increasing CioAB in the setting of high endogenous cyanide production.

## INTRODUCTION

Many bacteria have flexible and branched respiratory chains that allow them to obtain energy through aerobic and anaerobic respiration under diverse conditions (Kaila and Wikström, 2021). Terminal respiratory oxygen reductases are membrane-integrated enzymes that use an electron donor (cytochrome *c* or quinol) to catalyze the reduction of molecular oxygen to water at the end of aerobic respiratory chains. These enzymes belong to the evolutionary unrelated superfamilies of heme-copper oxidases (HCOs), alternative oxidases, and cytochrome *bd*-type oxygen reductases (cytbd) (Borisov et al., 2011; Borisov et al., 2021; Murali et al., 2021). While the HCO superfamily is widespread from bacteria to eukaryotic mitochondria, the cytbd superfamily is restricted to Bacteria and Archaea (Borisov et al., 2021; Murali et al., 2021).

Most redox reactions catalyzed by respiratory enzymes involve metal centers containing iron (as heme and iron-sulfur clusters) and copper. The HCO enzymes use a catalytic heme-copper binuclear center (BNC). In contrast, the cytbd enzymes contain heme but not copper ions (Borisov et al., 2021). The cytochrome *bd* oxygen reductases are quinol oxidases composed of two main integral membrane proteins, subunits I (CydA) and II (CydB), and an auxiliary small subunit (CydX) (cytochrome *bd*-I from *Escherichia coli*). CydA contains three heme groups (*b*558, *b*595 and *d*) and a quinol binding site. A hydrophobic region of CydA involved in quinol oxidation, called the Q-loop, has variable length. Based on this feature, cytbd reductases have been divided into two groups: L (long Q-loop) and S (short Q-loop). A third group includes cytochrome *bd* containing only b hemes, called cyanide insensitive terminal oxidases (CIO) (Cunningham et al., 1997; Borisov et al., 2011; Degli Esposti et al., 2015).

Cytochrome *bd* protects bacteria against several environmental or host-derived stressors, including nitrosative and oxidative stress and metal toxicity (Edwards et al., 2000; Mason et al., 2009; Borisov et al., 2013; Giuffrè et al., 2014; Al-Attar et al., 2016; Jones-Carson et al., 2016). Accordingly, in some bacteria, cytochrome *bd* mutants are severely compromised in virulence and intracellular viability (Small et al., 2013; Van Alst et al., 2022; Beebout et al., 2022). Cytochrome *bd* also protects bacteria from antibacterial drugs and are attractive targets for new antibiotics (Kaila et al., 2017; Beebout et al., 2021; Borisov et al., 2021; Kruth and Nett, 2023). The expression of cytochrome *bd* involves different regulatory systems. In *E. coli,* the *cydAB* genes reach maximum expression in microaerobic conditions by the concerted actions of ArcAB and Fnr, two oxygen-responsive regulatory systems (Fu et al., 1991; Tseng et al., 1996). In *Alishewanella sp.,* the *cydAB* genes are repressed by CydE, a GbsR-type regulator (Xia et al., 2018). The GbsR proteins comprise a poorly characterized subfamily of regulators belonging to the MarR (Multiple antibiotic resistance regulator) family of transcription factors. Genes encoding GbsR-like regulators are clustered in bacterial genomes with genes encoding cytochrome *bd* or transporters for osmoprotection (Ronzheimer et al., 2018; Xia et al., 2018).

*Chromobacterium violaceum* is a facultative anaerobic Gram-negative bacterium commonly isolated from soil and water that causes rare but deadly infections in humans and other animals (Yang and Li, 2011; Batista and da Silva Neto, 2017). Many *C. violaceum* traits such as production of violacein, exoenzymes, and cyanide are regulated by the LuxR-type quorum sensing (QS) system CviI/CviR (Mion et al., 2021; Batista et al., 2024). Although many species of the *Chromobacterium* genus are producers of cyanide (Mun et al., 2017; Short et al., 2018; Mion et al., 2021; Loo et al., 2023), it is unclear how these bacteria protect themselves from cyanide toxicity. *C. violaceum* has versatile respiratory capacity, exhibiting both cyanide-sensitive and -insensitive aerobic respiratory activities (Niven et al., 1975) and anaerobic nitrate respiration (Alves et al., 2021). Genes encoding both HCO and cytbd-type respiratory terminal oxidases are predicted in the *C. violaceum* genome, but their role and regulation remain unexplored. In this work, we demonstrated that a cytochrome *bd* (CioAB) repressed by a MarR-type transcription factor (CioR) protects *C. violaceum* against multiple inhibitors of HCO oxidases, including the endogenously produced poison cyanide.

## RESULTS

### Cytochrome *bd* encoded by the *cioRAB* operon protects *C. violaceum* from iron toxicity

We screened a *C. violaceum* transposon mutant library (Batista et al., 2024) for iron-susceptible transposon mutants on LB plates with 5 mM FeCl_3_ or 8 mM FeSO_4_. Mutant strains with impaired growth in excess iron had transposon insertions in genes related to regulatory systems and sugar metabolism (Table S1). One such mutant, with a transposon inserted in the gene CV_3659 (Table S1), was further investigated in this work. This gene, here named *cioR*, encodes a putative transcriptional regulator belonging to the GbsR subgroup of MarR family transcription factors (Ronzheimer et al., 2018; Xia et al., 2018). *cioR* is located adjacent to the genes CV_3658 and CV_3657, which encode the subunits I (CioA) and II (CioB) of a cytochrome *bd* quinol oxidase (Figure 1A). *C. violaceum* CioA presents a short Q-loop (Figure 1B) as in CioA from *Pseudomonas aeruginosa* (44.3% identity), CbdA from *Geobacillus stearothermophilus* (29.9% identity), and CydA from *Bacillus subtilis* (33% identity). RT-PCR reactions using RNA from WT (wild type) *C. violaceum* and primers corresponding to regions between the genes *cioR*-*cioA* and *cioR*-*cioB* amplified DNA bands with expected sizes (Figures 1A and C), indicating that these genes are co-transcribed, comprising the *cioRAB* operon.

**Figure 1.**
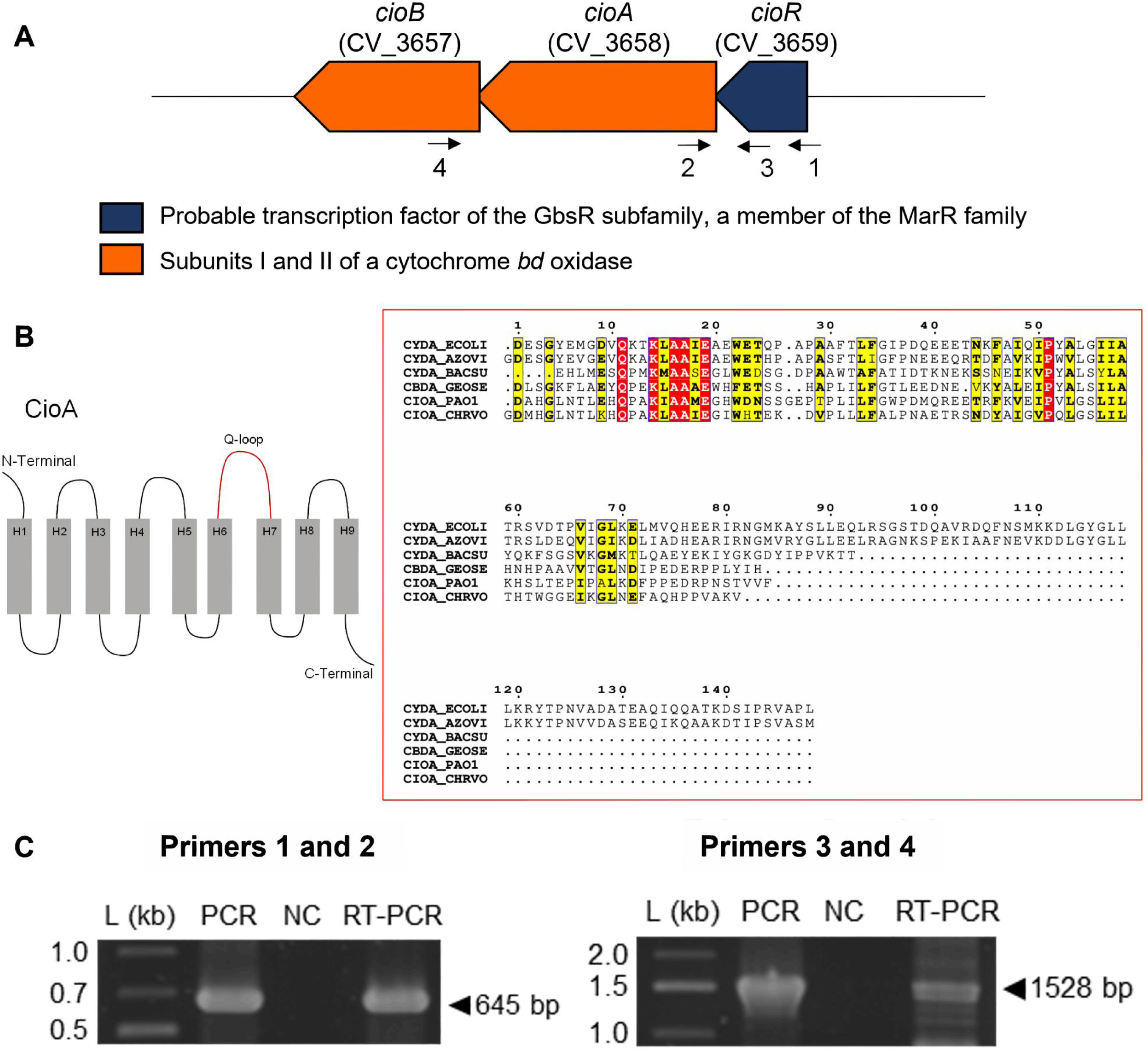
The *cioRAB* genes comprise an operon encoding a MarR-type regulator and a cytochrome *bd*. **A.** Genomic organization of the *cioRAB* genes in *C. violaceum*. Numbered arrows indicate primers used in RT-PCR. **B.** CioAB is a cytochrome *bd* with a short Q-loop in CioA. The diagram represents the membrane topology of CioA from *C. violaceum*. CioA has nine transmembrane segments (H1 to H9). Highlighted in red is the Q-loop involved in quinol oxidation. Multiple alignment of the Q-loop region from *C. violaceum* CioA with selected ortologs is shown. Proteins are from *C. violaceum* (CHRVO), *Escherichia coli* (ECOLI), *Azotobacter vinelandii* (AZOVI), *Bacillus subtilis* (BACSU), *Geobacillus stearothermophilus* (GEOSE), and *Pseudomonas aeruginosa* (PAO1). Identical and similar residues are boxed in red and yellow, respectively. **C.** Co-transcription of the *cioRAB* genes. The RT-PCR reactions amplified fragments of 645 bp (Primers 1 and 2) and 1528 bp (Primers 3 and 4) using RNA from WT *C. violaceum*. Conventional PCR was performed using genomic DNA (PCR) and RNA (NC), as controls. L, 1 Kb plus DNA Ladder (Thermo Scientific).

Since a *cioR* transposon mutant was impaired for growth at high iron concentrations (Table S1), we constructed null mutant strains to test the role of the *cioRAB* operon in iron tolerance (Figure 2). A Δ*cioR* mutant strain had no growth impairment in the iron susceptibility plate assays used in the transposon screen, whereas a Δ*cioAB* mutant failed to grow under the identical condition (Figure 2A). This suggested that the iron-susceptible phenotype of the *cioR* transposon mutant was due to a polar effect on the *cioAB* genes. Indeed, Δ*cioAB* but not Δ*cioR* strains exhibited increased sensitivity to iron and the iron-requiring antibiotic streptonigrin (SN) in growth curves (Figure 2B and C) and decreased production of siderophores on PSA-CAS plates (Figure 2D). All mutant strains grew similarly to WT under iron-deficient conditions (treatment with 2,2′-dipyridyl, DP) (Figure 2E). The sensitivity of a Δ*cioAB* mutant to FeCl_3_ and SN was further confirmed by survival and disk diffusion assays (Figure S1A and B). All of these iron-related phenotypes were abrogated by complementation of the Δ*cioAB* strain (Figure 2 and Figure S1A and B). Collectively, these data indicate that the cytochrome *bd* CioAB protects *C. violaceum* from iron toxicity. While our SN and siderophore assays suggest increased intracellular iron levels in a *C. violaceum* Δ*cioAB* strain, in *E. coli, cyd* mutants (*bd*-deficient) exhibit reduced iron content and increased siderophore production (Cook et al., 1998).

**Figure 2.**
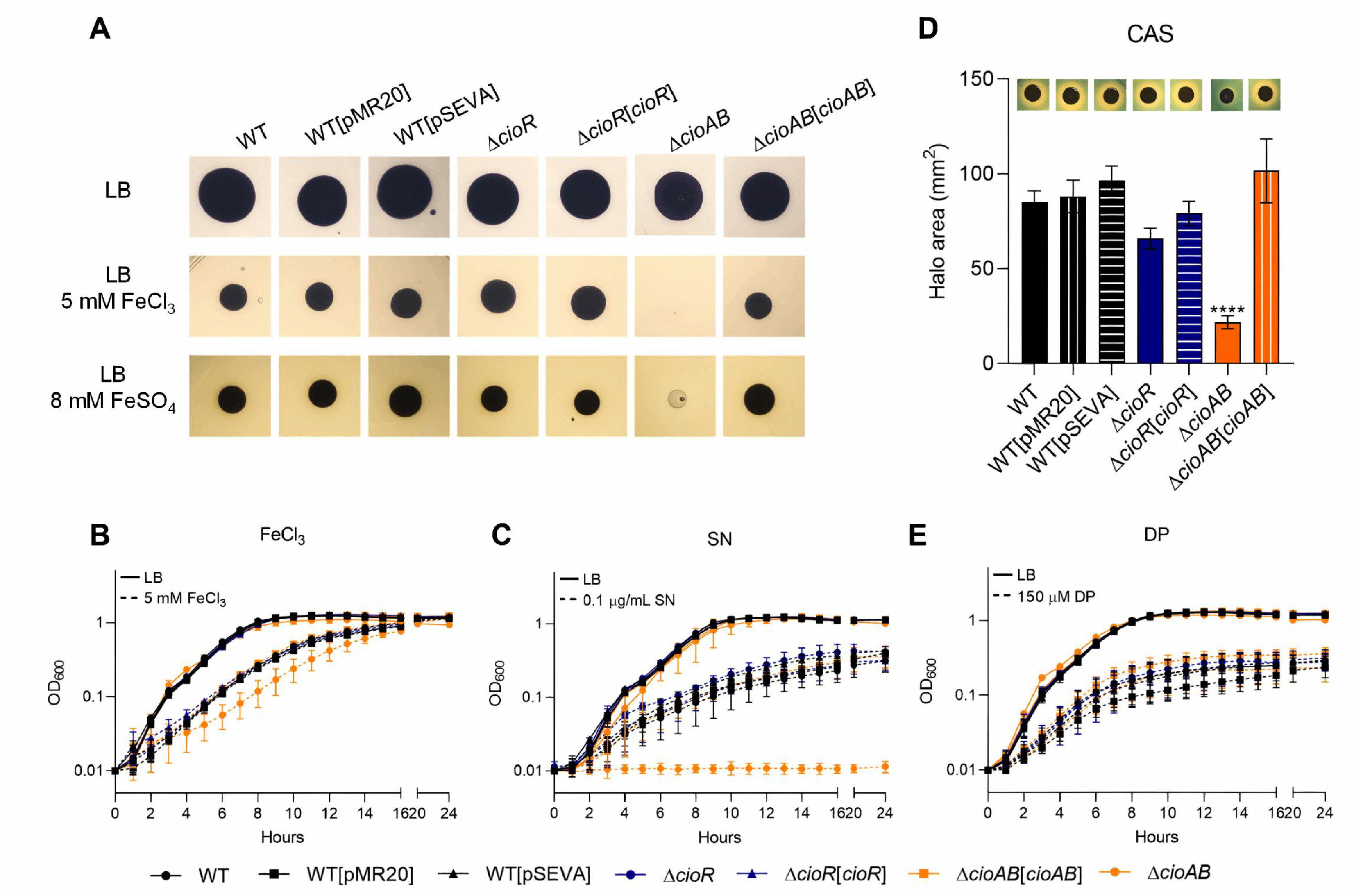
The cytochrome *bd* CioAB protects *C. violaceum* against iron toxicity. **A.** Iron susceptibility plate assays. The indicated strains were grown for 24 h at 37°C on LB plates without or with 5 mM FeCl_3_ or 8 mM FeSO_4_. **B, C, E.** Growth curves were obtained on a Bioscreen C microplate reader. The indicated strains were grown in LB medium in 96-well plates for 24 h at 37°C under agitation without (continuous lines) or with (dashed lines) the indicated treatments. **B.** 5 mM ferric chloride (FeCl_3_). **C.** 0.1 µg/mL streptonigrin (SN). **E.** 150 µM 2,2’-dipyridyl (DP). **D.** Mutation of *cioAB* leads to decreased siderophore halos. Measurement of the CAS halo diameter of the indicated strains. Data from three biological assays. Insert: illustrative PSA-CAS plates showing siderophore production (orange halos). ****p < 0.0001; when not indicated, not significant. One-way ANOVA followed by Tukey’s multiple-comparison test.

### The cytochrome *bd* CioAB confers resistance of *C. violaceum* to zinc, hydrogen peroxide, nitric oxide, sulfide and cyanide

The importance of the *cioRAB* operon in protecting *C. violaceum* under additional stress conditions was evaluated by growth curves comparing the WT strain with the *cio* mutants and complemented mutant strains (Figure 3). A Δ*cioAB* mutant was more susceptible to zinc (ZnCl_2_), hydrogen peroxide (H_2_O_2_), nitric oxide (sperNO), sulfide (cystine), potassium cyanide (KCN), and potassium ferrocyanide (K_4_[Fe(CN)_6_]·3H2O), compared with the WT and complemented strains (Figure 3). A Δ*cioR* mutant exhibited no growth defect under each of the tested stress conditions (Figure 3). Sensitivity of the Δ*cioAB* mutant to ZnCl_2_ and H_2_O_2_ was further confirmed by survival and disk diffusion assays (Figure S1A and B). These data indicate that *C. violaceum* relies on cytochrome *bd* CioAB to grow and survive in the presence of diverse toxic compounds. We hypothesized that the mechanism of stress resistance might involve the tolerance of CioAB to inhibition by these stressors, as many of them, including zinc, nitric oxide, sulfide, and cyanide, are known to inhibit HCO but not cytbd respiratory terminal oxidases in other bacteria (Cunningham et al., 1997; Mason et al., 2009; Chandrangsu and Helmann, 2016; Korshunov et al., 2016). The marked sensitivity of Δ*cioAB* to cyanide (Figure 3E) indicates that expression of CioAB is a key mechanism for *C. violaceum* resistance to this compound, which is endogenously produced by many *Chromobacterium* species (Mun et al., 2017; Short et al., 2018; Mion et al., 2021; Loo et al., 2023). WT *C. violaceum* was highly tolerant to cyanide, growing in concentrations up to mM of exogenously added potassium cyanide (Figure S1C). Curiously, when growth curves in LB were performed in large volumes (glass tubes instead of microplates), a Δ*cioAB* mutant reached a lower cell density after the late-exponential growth phase even in the absence of any exogenous stress (Figure S1D). This growth impairment of the Δ*cioAB* mutant in LB was also observed by survival assays of stationary growth phase cultures (Figure S1A). As cyanide accumulates from late-exponential growth phase in cultures of *C. violaceum* and other cyanogenic *Chromobacterium* species (Niven et al., 1975; Mun et al., 2017; Short et al., 2018; Loo et al., 2023), it is conceivable that the growth of a Δ*cioAB* mutant might be inhibited by endogenous cyanide.

**Figure 3.**
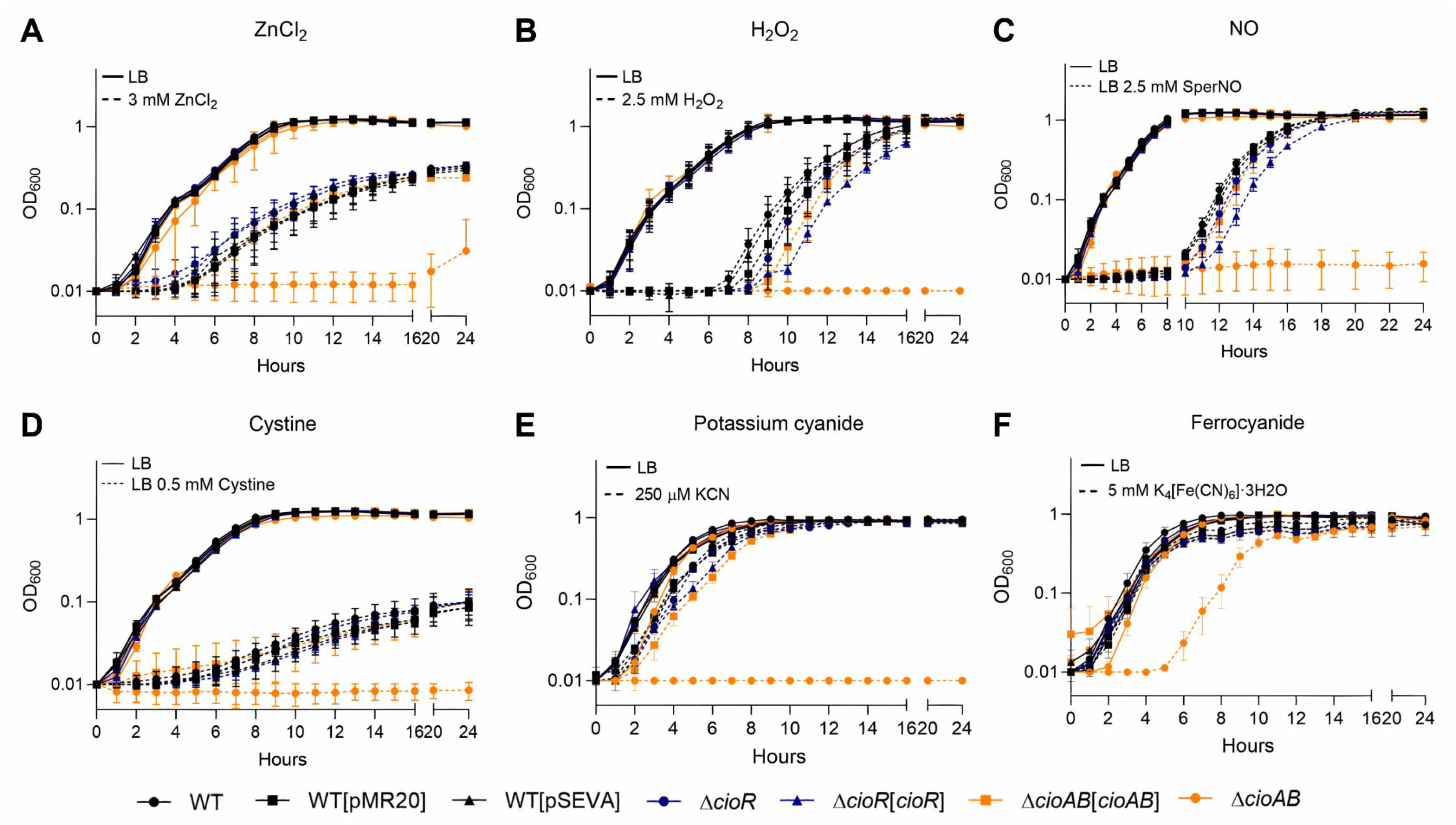
The cytochrome *bd* CioAB protects *C. violaceum* against toxicity of metal, oxidative, nitrosative, sulfide, and cyanide stresses. **A-G.** Growth curves were obtained on Bioscreen C or BioTek Epoch2 microplate readers. The indicated strains were grown in LB medium in 100-well or 96-well plates for 24 h at 37°C under agitation without (continuous lines) or with (dashed lines) the indicated treatments. **A.** 3 mM zinc chloride (ZnCl_2_). **B.** 2.5 mM hydrogen peroxide (H_2_O_2_). **C.** 2.5 mM spermine NONOate (SperNO, NO donor). **D.** 0.5 mM cystine (sulfide production). **E.** 250 µM potassium cyanide (KCN). **F.** 5 mM potassium ferrocyanide (K_4_[Fe(CN)_6_].3H_2_O).

### The *cioRAB* operon is repressed by CioR and activated by the quorum sensing regulator CviR

CioR is predicted to be a GbsR-type transcriptional regulator on the basis of sequence homology (Figures 1A). Because CioAB protects *C. violaceum* against cyanide (Figure 3E), and production of this compound relies on the QS system CviI/CviR (Mion et al., 2021; Loo et al., 2023), we tested the effect of QS and CioR on the expression of the *cioRAB* operon. The promoter region of the *cioRAB* operon was fused with the *lacZ* gene to measure its expression under low (LCD) and high (HCD) cell density, comparing expression between WT and mutant strains (Figure 4A). Beta-galactosidase assays with the WT strain showed that the *cioRAB* operon has a strong promoter that is upregulated from LCD to HCD. Mutation of *cioR* increased the expression of the *cioRAB* operon at LCD (Figure 4A). Expression of the *cioRAB* operon decreased in a Δ*cviR* mutant compared to WT under both cell density conditions (Figure 4A). These data indicate that the *cioRAB* operon is repressed by CioR at LCD and activated by CviR at LCD and HCD. To further demonstrate the role of CioR as a repressor of its own operon, we measured *cioA* and *cioB* transcripts in WT and Δ*cioR* strains by RT-qPCR. As expected, expression of both genes was increased in a Δ*cioR* mutant compared to WT and complemented strains (Figure 4B). To verify that CioR is a direct repressor of its own operon, we performed electrophoretic mobility shift assays (EMSA). The purified His-CioR protein was able to specifically bind to a DNA probe containing the promoter region of the *cioRAB* operon (Figure 4C), confirming its direct regulation by CioR.

**Figure 4.**
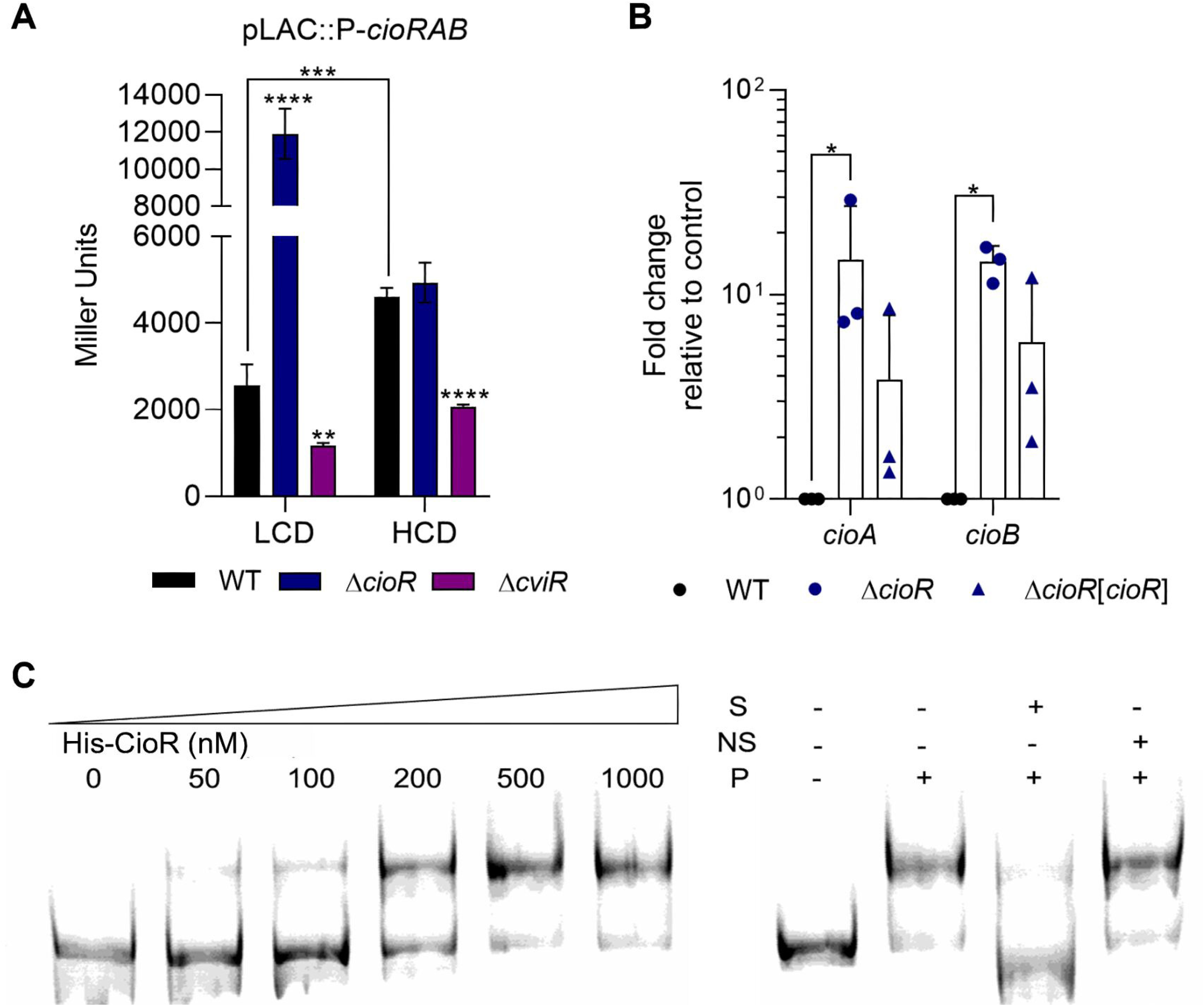
The *cioRAB* operon is directly repressed by CioR and activated by the QS regulator CviR. **A.** Expression of the *cioRAB* operon varies according to cell density and is regulated by CioR and CviR. Promoter activity was measured by β-galactosidase assays performed from WT, Δ*cioR*, and Δ*cviR* strains harboring the *cioRAB-lacZ* fusion. Strains were grown in LB medium in low cell density (LCD, OD_600_ 1.0) or high cell density (HCD, OD_600_ 4.0). Data are from five biological replicates. **p < 0.01; ***p < 0.001; ****p < 0.0001; when not indicated, not significant. Two-way ANOVA followed by Sidak’s multiple comparisons test. **B.** The *cioAB* genes are repressed by CioR. RNA obtained from WT, Δ*cioR*, and Δ*cioR*[*cioR*] strains grown in LB until OD_600_ 1.0 was reverse transcribed to cDNA and used for RT-qPCR reactions. Expression of *cioA* and *cioB* is shown as fold-change relative to the control condition (WT). Data are from three biological replicates. *p < 0.05; when not indicated, not significant. Two-way ANOVA followed by Tukey’s multiple-comparison test. **C.** CioR binds to the promoter region of the *cioRAB* operon. The indicated concentrations of His-CioR were used in EMSA assays with a *cioRAB*-FAM probe. Competition assays with unlabeled probes were performed to check binding specificity. S-Specific unlabeled probe; NS-Non-specific unlabeled probe; P-1 µM His-CioR protein.

### Expression of the *cioRAB* operon increases after exposure to exogenous cyanide

Because CioAB confers *C. violaceum* resistance to multiple stress conditions (Figure 2 and 3), we measured expression of the *cioRAB* operon in response to stressors, using the WT strain carrying a p*lacZ*::P*cioRAB* fusion (Figure 5). Eight distinct compounds were individually added to cultures of WT *C. violaceum* in LB at LCD for 30 min. Under these conditions, only cyanide caused a modest increase in the expression of the *cioRAB* operon. Addition of cyanide increased *cioRAB* expression two fold after 60-min exposure compared to an untreated LB control (Figure 5). These data indicate that, of the toxic compounds to which CioAB confers protection, only cyanide is an inducing signal for the *cioRAB* operon.

**Figure 5.**
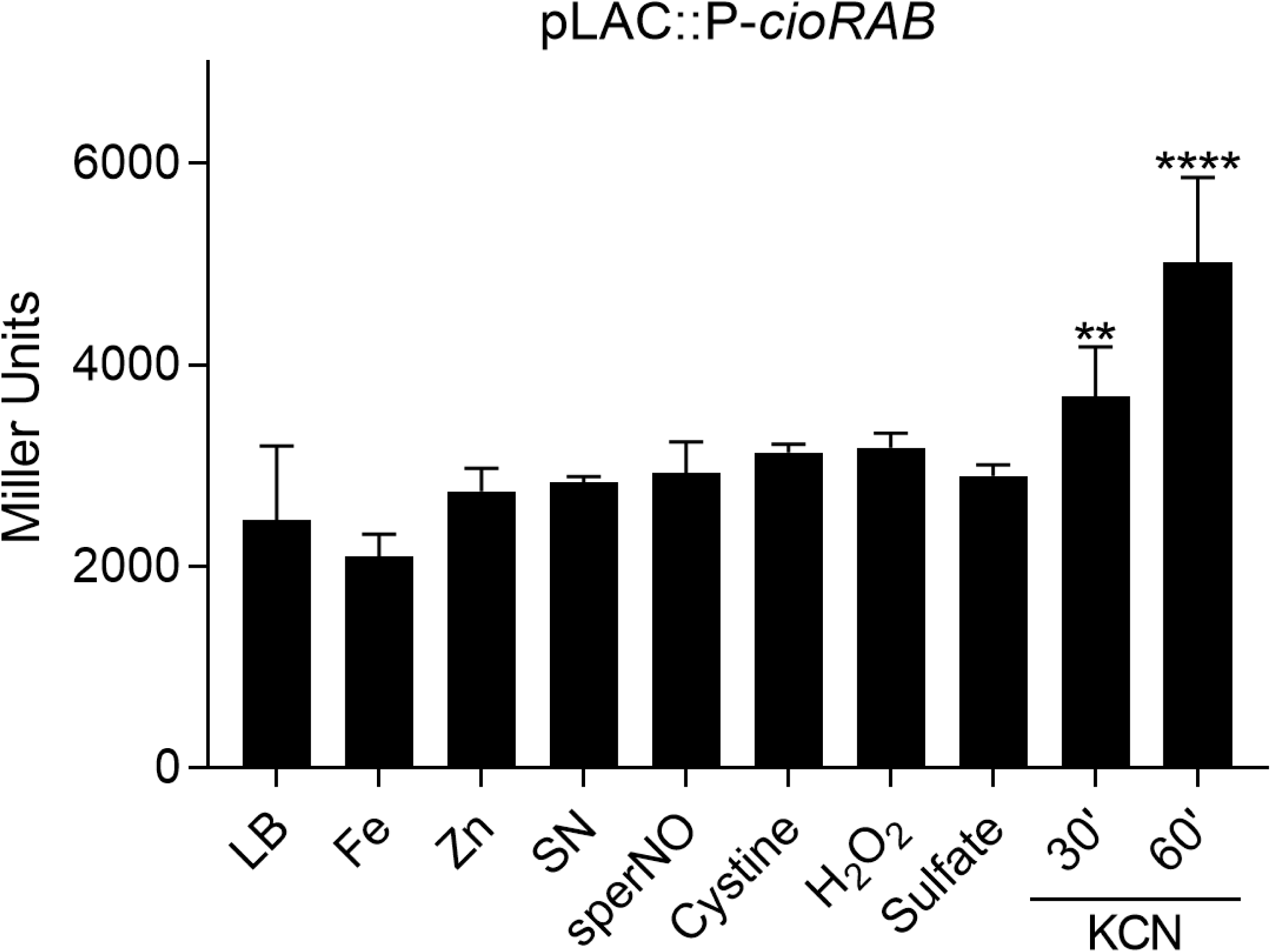
The *cioRAB* operon is induced in the presence of cyanide. Promoter activity of the *cioRAB* operon under treatment with different stress agents. For the β-galactosidase assays, cultures of the WT strain harboring the *cioRAB-lacZ* fusion were grown in LB medium until OD_600_ 0.5, and untreated or treated for 30 minutes with the following compounds: Fe, 5 mM FeCl_3_; Zn, 2 mM ZnCl_2_; SN, 0.5 µg/mL streptonigrin; sperNO, 1 mM spermine NONOate; Cystine, 0.5 mM cystine; H_2_O_2_, 10 mM hydrogen peroxide; Sulfate, 200 mM Na_2_SO_4_; Cyanide, 250 µM KCN for 30 and 60 minutes. Data are from three biological replicates. **p < 0.01; ***p < 0.001; when not indicated, not significant. Two-way ANOVA followed by Sidak’s multiple comparisons test.

## DISCUSSION

In this work, we demonstrated the direct regulation of the *cioRAB* operon by the GbsR-type regulator CioR and showed that a key role of the cytochrome *bd* CioAB is to protect *C. violaceum* against multiple environmental insults, including the endogenously produced poison cyanide and the host-derived antimicrobial molecule nitric oxide. Our findings further indicate that activation of the *cioRAB* operon by the quorum-sensing regulator CviR provide a regulatory mechanism to maximize CioAB expression during high endogenous cyanide production.

Our screen of a *C. violaceum* transposon mutant library revealed iron-susceptible mutants with transposon insertions in genes related to gene regulation and in a gene cluster for synthesis of polysaccharide precursors. A transposon screen in *Xanthomonas campestris* identified a β-(1-2)-glucan that protects against iron toxicity (Javvadi et al., 2018). A transposon insertion was found in *cioR*, but phenotypic characterization of Δ*cioR* and Δ*cioAB* null mutants revealed that deletion of *cioAB* but not *cioR* enhanced *C. violaceum* susceptibility to iron, zinc, hydrogen peroxide, nitric oxide, sulfide, and cyanide. The protective role of CioAB for many of these stressors is likely to involve tolerance of cytochrome *bd* to inhibition by compounds that are potent inhibitors of HCO terminal oxidases, including zinc (Chandrangsu and Helmann, 2016), sulfide (Korshunov et al., 2016), nitric oxide (NO) (Mason et al., 2009), and cyanide (Cunningham et al., 1997). Catalase activity of cytochrome *bd* accounts for hydrogen peroxide resistance (Borisov et al., 2013; Giuffrè et al., 2014; Borisov et al., 2015), and a rapid dissociation rate accounts for its insensitivity to nitric oxide (Mason et al., 2009). While cytochrome *bd* mutations result in iron deprivation in other bacteria (Cook et al., 1998; Edwards et al., 2020), a *C. violaceum* Δ*cioAB* mutant exhibited enhanced susceptibility to SN and decreased siderophore production, consistent with increased cell iron content. Previous work has shown that non-respiring *E. coli* accumulates NADH, which accelerates flavin reductase-mediated electron transfer to free flavins, which in turn can reduce free iron to catalyze the Fenton reaction and promote DNA damage (Woodmansee and Imlay, 2002). As SN depends on Fe(II) for its killing activity (Gupta and Imlay, 2023), we suggest that a similar mechanism could account for the iron, H_2_O_2_, and SN susceptibility of a *C. violaceum* Δ*cioAB*. However, an explanation for the apparently divergent effects of cytochrome *bd* on iron content and siderophore production in *C. violaceum* is not presently known.

In this work, we have identified different conditions and transcriptional regulators controlling the expression of the *cioRAB* operon in *C. violaceum*. We found that CioR, belonging to the GbsR subfamily of MarR family transcription factors (Ronzheimer et al., 2018; Xia et al., 2018), acts as a direct repressor of its own operon. The only previously characterized GbsR-type regulator associated with cytochrome *bd* genes, CydE, represses the *cydEAB* operon of *Alishewanella* and appears to respond to sulfate (Xia et al., 2018). In *C. violaceum,* we found that cyanide but not sulfate induces the *cioRAB* operon. It remains to be determined whether CioR is a cyanide-sensing regulator, like MpaR in *P. aeruginosa* (Smiley et al., 2024), or senses other molecules, similarly to TstR, which senses sulfite and Fe(III) to control cyanide resistance in *Lactobacillus brevis* (Pagliai et al., 2014). We have previously demonstrated that the MarR family regulators OhrR and OsbR coordinate responses against oxidative stress in *C. violaceum* (da Silva Neto et al., 2012; Previato-Mello et al., 2017; Alves et al., 2021). OsbR has a large regulon, including the *nar* genes involved in anaerobic nitrate respiration and the *cioRAB* operon (Alves et al., 2021). Interestingly, in addition to direct repression by OsbR (Alves et al., 2021) and CioR, we have shown here that the *cioRAB* operon is also regulated by cell density and activated by the QS regulator CviR, a result that is consistent with a recent global transcriptome analysis (Batista et al., 2024). As production of cyanide is also a QS-regulated process (Mion et al., 2021; Loo et al., 2023) employed by *Chromobacterium* species for predation evasion (Mun et al., 2017), bacterial competition (Loo et al., 2023), and larval killing (Short et al., 2018), our data demonstrating the marked susceptibility of a Δ*cioRAB* mutant to cyanide suggest that cyanide tolerance conferred by the cytochrome *bd* CioAB is likely to play an important role in many facets of the *C. violaceum* biology. In *P. aeruginosa*, QS and cyanide regulation of *cioAB* contributes to cooperative behavior in bacterial populations (Yan et al., 2019).

Many bacterial pathogens require cytochrome *bd* enzymes during host infection, including *Vibrio cholerae* (Van Alst et al., 2022), *Salmonella enterica* serovar Typhimurium (Jones-Carson et al., 2016), *Mycobacterium tuberculosis* (Cai et al., 2021), uropathogenic *Escherichia coli* (Beebout et al., 2022), and *Listeria monocytogenes* (Corbett et al., 2017). In some cases, the role in virulence involves cytochrome *bd*-mediated protection against nitric oxide (NO) released by host cells (Jones-Carson et al., 2016; Beebout et al., 2022). Recently, macrophage-produced NO has been implicated in controlling *C. violaceum* infection in the mouse liver within granulomas (Harvest et al 2023). Additional studies will be required to elucidate the specific role of the cytochrome *bd* CioAB in *C. violaceum* virulence, given its protective role against multiple stress conditions including cyanide and nitrosative stress.

## MATERIALS AND METHODS

### Bacterial strains, plasmids, and growth conditions

Bacterial strains and plasmids used in this work are described in Table 1. *E. coli* and *C. violaceum* strains were cultured in Luria-Bertani (LB) medium. When appropriate, cultures were supplemented with kanamycin (50 μg/mL), ampicillin (100 μg/mL), or tetracycline (10 ug/mL).

**Table 1.**
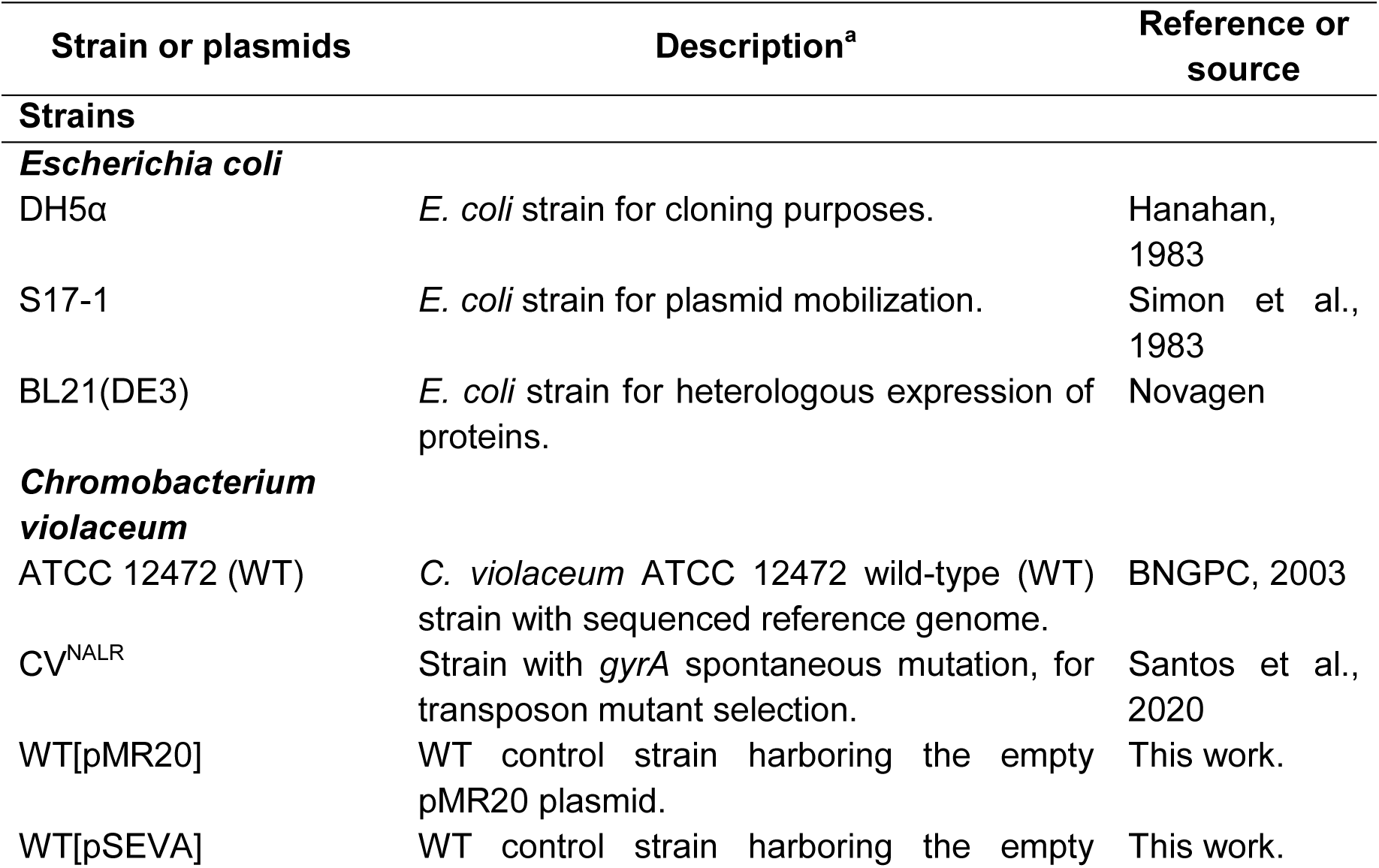
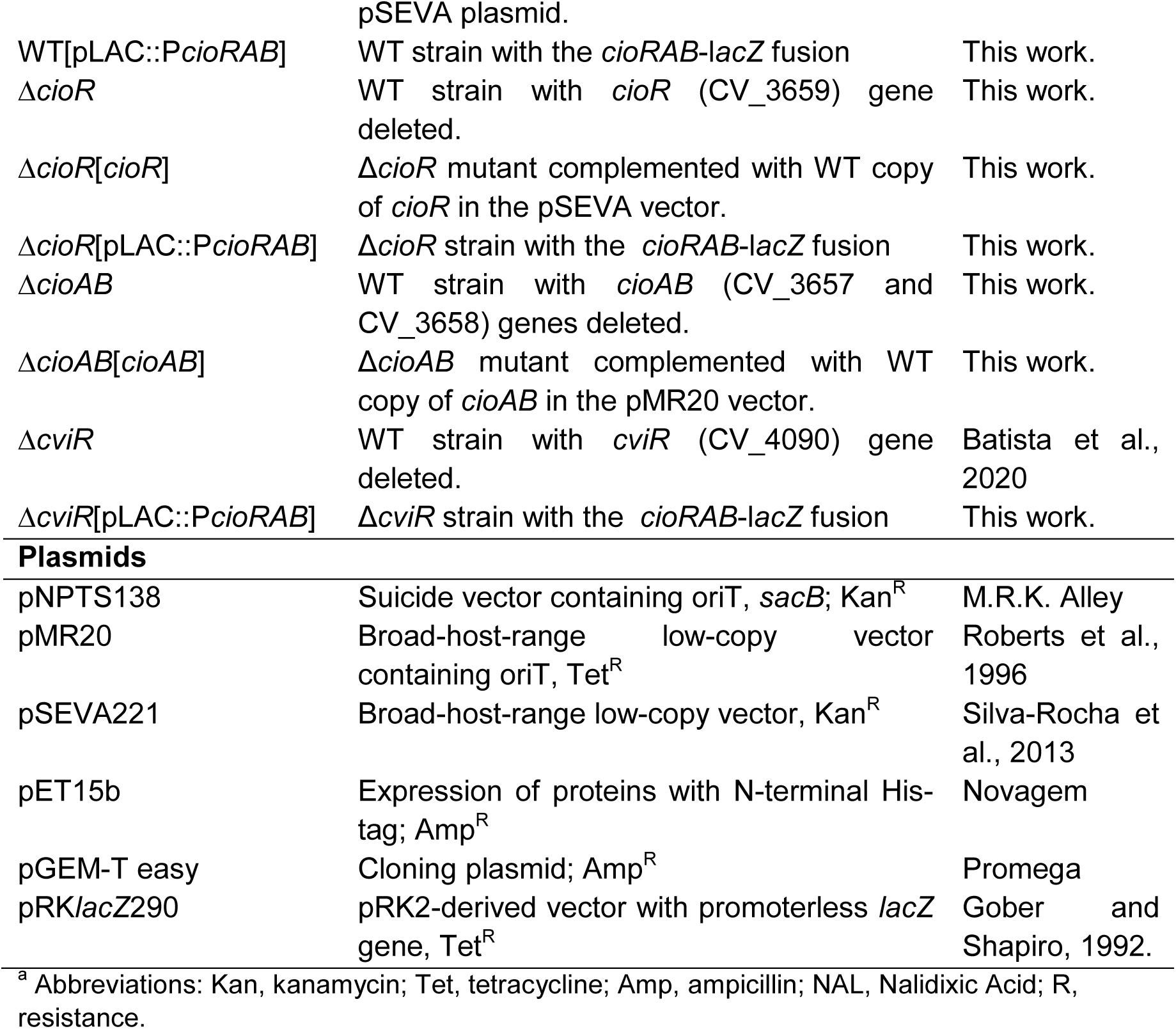
Strains and Plasmids.

### *In silico* analysis of CioA

Protein sequences of cytochrome *bd* subunit I from *C. violaceum* (CioA), *E. coli* (CydA), *A. vinelandii* (CydA)*, B. subtilis* (CydA), *G. stearothermophilus* (CbdA), and *P. aeruginosa* (CioA) were retrieved from Uniprot (https://www.uniprot.org/). Transmembrane helices were predicted using TMHMM - 2.0 (https://services.healthtech.dtu.dk/services/TMHMM-2.0/).

Amino acids of the Q-loop region (located between the transmembrane helices six and seven) were selected for multiple sequence alignment using Clustal Omega (https://www.ebi.ac.uk/Tools/msa/clustalo/) and visualization with ESPript 3.x (https://espript.ibcp.fr/ESPript/ESPript/).

### Screening of a transposon mutant library for iron toxicity

A previously obtained *C. violaceum* 10,000-mutant library (Batista et al., 2024) was screened for iron susceptibility by spotting 10 µL of overnight cultures of individual clones on LB plates supplemented with 5 mM FeCl_3_ or 8 mM FeSO_4_. Transposon mutant strains with growth impairment after incubation for 24 hours at 37°C under iron excess were selected for identification of transposon insertion sites by semi-degenerate PCR (Table S2), followed by Sanger sequencing as previously described (Jacobs et al., 2003; Santos et al., 2020; Batista et al., 2024). Null *cio* mutants were also tested in the iron-supplemented plate assays.

### Construction of *C. violaceum* mutant and complemented strains

All null-mutant strains were derived from the *C. violaceum* wild-type (WT) strain ATCC 12472. In-frame null deletions of *cioR* (Δ*cioR*) and *cioAB* (Δ*cioAB*) were generated by allelic exchange mutagenesis using sucrose for counter-selection, as previously described (da Silva Neto et al., 2012; Batista et al., 2019; Santos et al., 2020; Batista et al., 2024). Null mutant strains were complemented by providing wild-type copies of the mutated genes cloned into plasmids pMR20 (for *cioAB*) or pSEVA (for *cioR*). Primers used for cloning, sequencing and confirmation of mutant and complemented strains are listed in Table S2.

### Growth curves

*C. violaceum* wild-type, mutant, and complemented strains were grown in LB medium overnight. Cultures were diluted to an optical density at 600 nm (OD_600_) of 0.01 in LB that was untreated or treated with individual stress agents to a final volume of 150 µL in 96-well or 300 µL in 100-well microplates. Cultures were grown aerobically with shaking at 37°C in Bioscreen C or BioTek Epoch2 microplate readers. Growth was monitored by measuring OD_600_ every 60 min. Experiments were performed with six replicates. Treatment conditions are indicated in the respective figures. A set of growth curves in large volumes (5 mL cultures) were obtained in glass tubes containing LB that were incubated at 37°C in agitation. Aliquots were taken to measure OD_600_ in an Eppendorf BioPhotometer.

### Survival assays

*C. violaceum* wild-type, mutant or complemented strains were grown in LB overnight. Cultures were diluted to OD_600_ 0.01 in LB and then grown aerobically with shaking at 37°C until OD_600_ reached 0.5. Cultures were untreated or treated with stress agents and incubated for 20 h with agitation at 37°C. Serial dilutions were performed in phosphate-buffered saline (PBS) and plated on LB for colony-forming unit (CFU) enumeration after 24 h of incubation. Experiments were performed with six replicates. Treatment conditions are indicated in the respective figures.

### Disk diffusion assays

*C. violaceum* wild-type, mutant, and complemented strains grown overnight in LB were diluted to OD_600_ 1.0 in LB. Twenty μL of each culture were embedded in 20 mL of LB agar. Wells were created on the plate, and 30 μL aliquots of 10 μg/mL SN, 10 mM H_2_O_2_, 80 mM ZnCl_2_, or 100 mM FeCl_3_ were applied to individual wells. After incubation for 24 h at 37°C, halos of growth inhibition around each well were measured. Halo diameter was quantified using ImageJ software. Experiments were performed with three biological replicates.

### Siderophore production assay

Siderophores were detected using the universal chrome azurol S (CAS) agar plate assay (Schwyn and Neilands, 1987) modified by replacing MM9 medium with peptone-sucrose agar (PSA) (Batista et al., 2019; Batista et al., 2024). Ten microliters of *C. violaceum* cultures were spotted onto PSA-CAS agar plates, and siderophore production was evaluated by detection of orange halos that appeared after incubation for 24 hours at 37°C.

### Co-Transcription by RT-PCR

The *C. violaceum* wild-type strain was grown in LB to OD_600_ 4.0. Total RNA was extracted using Trizol reagent (Invitrogen) and purified with Direct-zol™ RNA Miniprep Plus (Zymo Research). RT-PCR was performed with the SuperScript III One-Step RT-PCR System with Platinum Taq High Fidelity DNA Polymerase (Invitrogen). One microgram of each RNA sample and specific primers (Table S2) that amplify regions from *cioR* to *cioA* (645 bp) and *cioR* to *cioB* (1528 bp) were used in the reactions. PCR using Taq DNA polymerase and the same primer sets, was performed with genomic DNA (positive control) and RNA (negative control) as templates.

### Gene Expression by RT-qPCR

*C. violaceum* wild type, Δ*cioR*, and Δ*cioR* complemented strains were grown in LB to OD_600_ 1.0. Total RNA was extracted using TRIzol according to the manufacturer’s protocols. Five hundred nanograms of total RNA from each sample were converted to cDNA using the QuantiTect reverse transcription kit (Qiagen). Quantitative PCR (qPCR) reactions were performed and quantified in a Bio-Rad CFX96 (Bio-Rad), using SYBR Green master mix, specific primers (Table S2), and 2 μL of cDNA. Data from three biological replicates were normalized by an endogenous control (gene CV_3376) and a reference condition (WT in LB).

### Construction of transcriptional *lacZ* fusions and β-galactosidase assay

The region upstream of the *cioRAB* operon was amplified by PCR with designated primers (Table S2), cloned into the pGEM-T easy plasmid (Promega), and subcloned into the pRK*lacZ*290 vector to generate a transcriptional fusion to the *lacZ* gene. This construct was introduced into *C. violaceum* WT, Δ*cioR*, and Δ*cviR* strains. These strains were cultured in LB to OD_600_ 1.0 (LCD, low cellular density) or OD_600_ 4.0 (HCD, high cellular density). For stress conditions, the WT strain was grown in LB to OD_600_ 0.5, and cultures were untreated or treated for 30 minutes with stress agents as indicated in the respective figures. All cultures were assayed for β-galactosidase activity using a previously described protocol modified for *C. violaceum* (Santos et al., 2020; Batista et al., 2024).

### Expression and purification of CioR

The coding region of *cioR* was PCR-amplified (Table S2) and cloned into the pET15b vector. The recombinant histidine-tagged protein (His-CioR) was overexpressed in *E. coli* BL21(DE3) by induction with 1 mM isopropyl-D-thiogalactopyranoside (IPTG) for 2 h at 37°C in LB. The His-CioR protein was purified from the soluble cell fraction using a 5 mL HisTrap HP column (Cytiva Life Sciences) on an AKTA Explorer FPLC system (Cytiva Life Sciences). Elution samples were concentrated and desalted by dialysis in storage buffer (20 mM Tris (pH 7.6), 150 mM NaCl, 0.1 mM EDTA, 5 mM DTT). Protein purification was checked by 15% SDS-PAGE.

### Electrophoretic mobility shift assay (EMSA)

A DNA 6-FAM labeled probe containing the promoter region of the *cioRAB* operon was amplified by PCR (Table S2). DNA binding reactions were performed in interaction buffer (20 mM Tris-HCl [pH 8.0], 50 mM NaCl, 1.5 mM MgSO_4_, 0.5 mM CaCl_2_, 0,1 mg/mL bovine serum albumin, 1 mM DTT, 0.05% NP-40, 10% glycerol), 0.1 mg/mL competitor salmon sperm DNA, 50 ng labeled 6-FAM DNA probe and different concentrations of His-CioR in a final volume of 20 μL. All interaction reactions were incubated at 37°C for 25 min. Samples were separated by native 5% polyacrylamide gel electrophoresis in Tris-borate (TB) buffer. Competition assays were performed using 1 µM of His-CioR in the presence of a 10-fold excess of unlabeled specific (*cioRAB* promoter region) or non-specific (CV_3376 coding region) probes. The fluorescence signal was captured by the Azure Sapphire Biomolecular Imager.

## ACKNOWLEDGMENTS

This research was supported by grants from the São Paulo Research Foundation (FAPESP; grants 2018/01388-6 and 2021/06894-0) and Fundação de Apoio ao Ensino, Pesquisa e Assistência do Hospital das Clínicas da FMRP-USP (FAEPA). During the course of this work, B.B.B. (grants 2018/19058-2 and 2021/09170-2) and V.M.L. (grant 2020/15268-2) were supported by fellowships from FAPESP and CAPES (Coordenação de Aperfeiçoamento de Pessoal de Nível Superior). J.F.S.N. is Research Fellow from CNPq (Conselho Nacional de Desenvolvimento Científico). W.R.W. and F.C.F. are supported by the U.S. National Institutes of Health (AI 150041).

---

**Supplementary Figure 1. Phenotypic tests of *C. violaceum* WT and *cio* mutants under various treatment conditions. A and B.** The Δ*cioAB* mutant is susceptible to multiple stress conditions. **A.** Survival assays. Indicated strains were grown in LB until OD_600_ 0.5. The cultures were untreated or treated and incubated for 20 h under agitation at 37°C. Serial dilutions were plated on LB for CFU counting after 24 h of incubation. **B.** Disk diffusion assays. The strains were embedded in LB plates and 30 µL aliquots of various compounds were added to wells. After incubation at 37°C for 24 h, halos of growth inhibition were measured. Data are from three biological replicates. ****p < 0.0001; ***p < 0.001; **p < 0.01; *p < 0.05; when not indicated, not significant. Significance was determined by two-way ANOVA followed by Tukey’s multiple-comparison test. **C.** Susceptibility of *C. violaceum* to cyanide. Growth curves were obtained on a BioTek Epoch2 microplate reader. The wild-type strain was grown in LB without or with different concentrations of potassium cyanide in 96-well plates for 24 h at 37°C under agitation. **D.** Cultures of the Δ*cioAB* mutant reached a lower cell density after late-exponential growth phase. Growth curves were obtained by measurement of OD_600_ of the cultures during the first eight hours (intervals of 1 h) and at 24 h. Indicated strains were grown in LB medium in glass tubes at 37°C under agitation.

